# Computational discovery of co-expressed antigens as dual targeting candidates for cancer therapy through bulk, single-cell, and spatial transcriptomics

**DOI:** 10.1101/2023.10.17.562711

**Authors:** Evgenii Chekalin, Shreya Paithankar, Rama Shankar, Jing Xing, Wenfeng Xu, Bin Chen

## Abstract

**Motivation:** Bispecific antibodies (bsAbs) that bind to two distinct surface antigens on cancer cells are emerging as an appealing therapeutic strategy in cancer immunotherapy. However, considering the vast number of surface proteins, experimental identification of potential antigen pairs that are selectively expressed in cancer cells and not in normal cells is both costly and time-consuming. Recent studies have utilized large bulk RNA-seq databases to propose bispecific targets for various cancers. But, co-expressed pairs derived from bulk RNA-seq do not necessarily indicate true co-expression of both markers in the same cell. Single-cell RNA-seq (scRNA-seq) can circumvent this issue but the issues in dropouts and low-coverage of transcripts impede the large-scale characterization of co-expressed pairs.

**Results:** We present a computational pipeline for bsAbs identification which combines the advantages of bulk and scRNA-seq while minimizing the issues associated with using these approaches separately. We select Hepatocellular Carcinoma (HCC) as a case study to demonstrate the utility of the approach. First, using the bulk RNA-seq samples in the OCTAD database, we identified target pairs that most distinctly differentiate tumors cases from healthy controls. Next, we confirmed our findings on the scRNA-seq database comprising 39,361 healthy cells from vital organs and 18,000 malignant cells from HCC. The top pair was GPC3∼MUC13, where both genes are co-expressed on the surface of over 30% of malignant HCCs and have very low expression in other cells. Finally, we leveraged the emerging spatial transcriptomic to validate the co-expressed pair *in situ*.

**Availability and Implementation:** A standalone R package for bsAbs identification in bulk data is available via GitHub (https://github.com/Lionir/bsAbsFinder).

## INTRODUCTION

Bispecific antibodies (bsAbs) have garnered much interest in cancer therapeutic discovery recently. Several types of bsAbs have been explored based on the types of biological targets and modes of action. The majority of bsAbs under clinical investigation are bispecific immune cell engagers where one arm of the bsAbs targets an established immune cell antigen and another arm links to a tumor cell antigen. For instance, Blinatumomab targets CD3 antigen of T cells and CD19 surface antigen of B cells for the treatment of B-cell malignancies [1]. Another less-studied bsAbs class aims to target two tumor-associated antigens such that the drug could improve selectivity towards tumor cells, while minimizing side effects in normal tissues, or modulate two functional pathways in the tumor to overcome treatment resistance [1,2]. Similarly, Chimeric antigen receptor (CAR) T-cell that simultaneously target two tumor-associated antigens can also enhance antitumor activity and circumvent escape mechanisms in solid tumors [3,4]. Recent advances in protein engineering allow designing protein logic to target cells with precise combinations of surface antigens [5,6], yet the optimal combinations of antigens remain largely unexplored. Considering the wide range of surface antigens yielding millions of possible bsAbs pairs to evaluate, computational identification of bsAbs pairs that are expressed only in cancer cells but not in normal cells has promising application in bsAbs-based cancer immunotherapy.

As rich bulk RNA-seq data has been generated for various cancers and normal tissues, it has been extensively reused to pinpoint therapeutic targets and biomarkers. A recently proposed computational approach employed Boolean logic to characterize highly co-expressed antigen pairs in tumor vs. healthy samples in bulk RNA-seq databases. They utilized large bulk RNA-seq databases such as The Cancer Genome Atlas Program (TCGA, https://www.cancer.gov/tcga) and Genotype-Tissue Expression portal (GTEx) as sources of tumors and healthy control to identify novel CAR T-cells marker candidates [7]. However, heterogenous cell population of bulk RNA-seq limits its potential to discern the precise expression of targets at the individual cell level within cancer tissues, potentially leading to a significant number of false positive antigen pairs. Whereas, scRNA-seq offers a high-resolution quantification of target expression in individual cells, but the challenges in addressing the dropout issue and accurate identification of cell types restrict its direct application in bsAbs discovery.

In this study, we introduce a computational pipeline for identifying bsAbs that leverages the strengths of bulk and scRNA-seq while mitigating the challenges typically associated with using these methods in isolation. We first harnessed high-quality bulk RNA-seq data compiled from multiple open databases to identify novel pairs. We developed computational approaches to find the pairs with a similar high expression level within tumor samples (case) and a low or zero expression level within non-tumor samples (control). Along with normal and/or adjacent normal tissues related to the given cancer, normal vital organs were also included in the control dataset. The resulted pairs from bulk RNA-seq analysis were subsequently verified in an assembled scRNA-seq dataset to confirm their co-expression in the same cell type (Fig. 1A,B). Further, spatial transcriptomic data was used for external validation to visualize the co-expression pattern of the top pair in cancerous and healthy tissues. The utility of proposed pipeline was demonstrated in Hepatocellular Carcinoma (HCC) bsAbs discovery.

**Figure 1:**
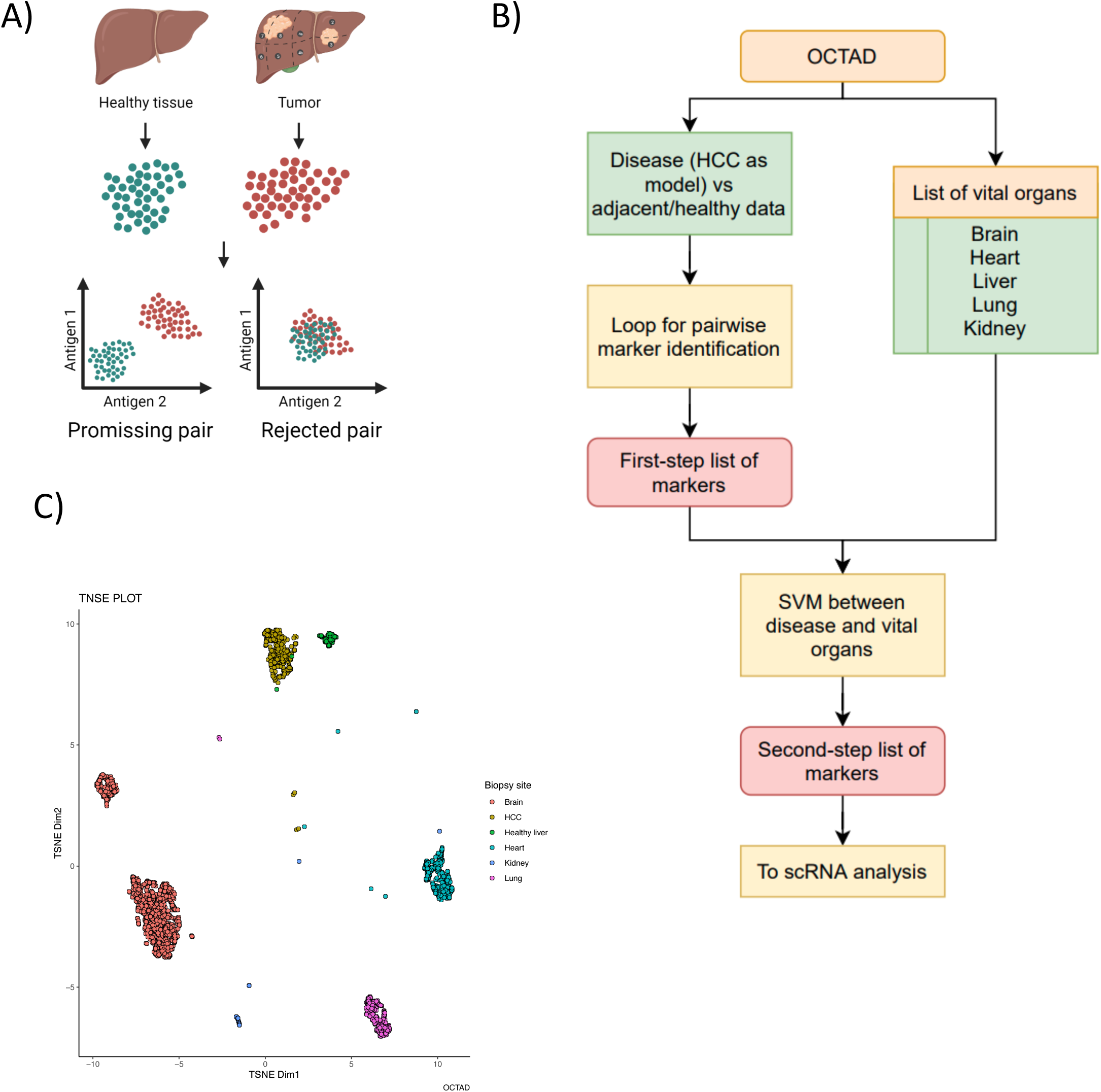
Identification of bulk RNA-seq-based bsAbs pairs**. A:** Schematic of the bulk RNA-seq based bsAbs pair selection,B: Workflow of bsAbs pair selection, **C**: t-distributed stochastic neighbor embedding (t-SNE) plot of the OCTAD data for HCC, normal liver, and normal vital organs used for the verification of the non-specific cytotoxicity of the proposed bsAbs candidate pairs.

## MATERIALS AND METHODS

### A. Data collection and preprocessing

#### Bulk RNA-Seq Database

We utilized the OCTAD database, a collection of databases including TCGA, GTEX, and MET500 [8]. It contains a total of 19,127 samples including 11,715 samples from 130 distinct cancers and 1,950 healthy samples from 5 vital organs (1,148 brain samples from various regions, 376 heart samples either from the left ventricle or atrial appendage, 28 samples from the kidney cortex, 110 liver samples, and 288 lung samples) that were processed from raw sequence data under the same pipeline [9]. Recent studies demonstrated the feasibility of performing integrative analysis of these samples in various tasks [10–12]. For current study, 369 HCC case samples were compared against all healthy control samples (Supplementary Table S1, S2).

#### Surface Protein Database

The COMPARTMENTS [13] database provides protein subcellular localization evidence built by integrating multiple sources like experimental datasets, sequence-based predictions, and manual literature curation, or by automated text-mining. Subcellular compartment terms were mapped to the Cellular Component subset of the Gene Ontology. Data from disparate sources were standardized by assigning confidence scores for all protein-compartment associations. We selected only 3,806 surface protein-coding genes from this database based on confidence score > 4 (Supplementary Table S3).

#### Single-cell RNA-seq Data

For the single cell data of the case samples, we used 70,000 cells of GEO dataset GSE149614 [14] which were collected from 10 HCC patients. For control dataset, we merged nearly 40,000 cells (Supplementary Table S4) from five distinct sources: 10,372 cells from the liver [15], 10,360 cells from the lung [16], 8,710 cells from the kidney nephrectomy [17], 6,134 cells from the brain cortex [18] and 3,785 cells from the left atrium and ventricle of normal heart [19]. We performed quality control (QC), normalization, and batch correction of the scRNA-seq datasets using Seurat [20], Monocle [21], and Harmony [22]. We used Seurat to process and normalize the data, and then filtered out cells that expressed fewer than 200 genes or those with mitochondrial genes exceeding 5%. The remaining datasets underwent log-normalization with a 10k scale factor. The top 2000 variable features were selected for cell type labeling. The SingleR R package [23] which includes human primary cell atlas dataset [24] as a reference was used to classify cells. The human cell atlas contains the expression of over 40 cell types, including liver cells.

#### Spatial Transcriptome Data

Spatial transcriptomics (ST) is an emerging technology for profiling gene expression at the spatial resolution. We downloaded the HCC spatial transcriptome data from http://lifeome.net/supp/livercancer-st/data.htm [25]. The ST tissue samples were processed using the SpaceRanger (10X Genomics) by the original authors.

### B. Data Analysis

#### Identification of bsAbs pairs based on surface marker expression in bulk RNA-seq

High sensitivity and high specificity are two essential attributes for a candidate marker. High sensitivity indicates high expression rates in malignant tumor cells, while high specificity means that a great proportion of non-malignant cells do not express the marker. A qualified marker should be highly expressed in malignant cells and also should have low or zero expression in non-malignant cells in the same tissue. Moreover, it should exhibit low or no expression in vital organs such as the brain, liver, lung, kidney, and heart to minimize therapy-related toxicity.

We aim to identify bsAbs pairs not only based on their higher expression in case samples but also how they maximally differentiate between case vs control samples. The pairwise combination of 3,806 genes from COMPARTMENT database has yielded over 7 million distinct bsAbs pairs. We used the following formula to identify promising bsAbs pairs:

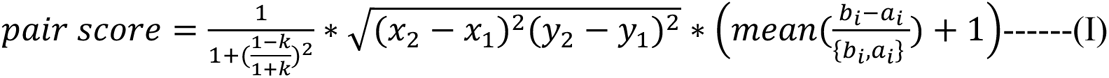

Where k is the slope of the linear regression defined by medoids of case (tumor) and control (non-tumor) cluster; (x_1_,y_1_) are the coordinates of the medoid for the control cluster, (x_2_,y_2_) are the coordinates of the medoid for the case cluster and 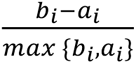 defines the silhouette of the case and control cluster.

Note that the resulting score reflected only marker candidacy regardless of the group separated by markers. Therefore, to obtain the pairs for the case cluster, we also used the Boolean approach: both medoid coordinates for the case cluster should be greater than those for the control cluster, meaning both genes of the marker pair should exhibit high expression in the case cluster and low or negligible expression in the control cluster. Since the distribution of the scores was beta-like, to obtain p-values we used a permutation test with FDR correction. Next to ensure higher specificity, we checked the expression of marker genes of significant bsAbs pairs in OCTAD reference normal vital organ samples (Supplementary Table S2).

To compute the sensitivity and specificity of candidate bsAbs pairs, we used SVM (Support Vector Machine) available in R package e1071 along with the ROCR R package. We performed two separate sensitivity and specificity comparison between dual markers (bsAbs) and single markers. Since the whole bsAbs identification task can be viewed as a 2-feature classification problem, we trained two distinct SVM models: one for the case vs. adjacent normal tissue and another for case vs. vital organs. We computed the mean delta between the sensitivity and specificity of both models (i.e., bsAbs pairs and individual markers in the pairs).

#### Verification of cell-specific co-expression of the bsAbs pairs using scRNA-seq data

To confirm the bsAbs identified through bulk RNA-seq, we utilized the single-cell RNA-seq data. As tumor is a mixture of immune cells, stromal cells, and hepatocytes, with the latter including malignant and non-malignant hepatocytes, it was important to concentrate only on the malignant cells in the tumor while computing true co-expression percentage. The inclusion of non-malignant parenchymal cells in bulk tumor tissues could have led to the identification of false-positive markers. To distinguish malignant cells from non-malignant ones, we turned to the inferCNV package [26,27] with default parameters and referenced healthy hepatocytes obtained from the human liver cell atlas dataset [15]. After selecting the malignant cells from the scRNA-seq dataset, we validated the list of markers identified at the tissue level and chose those expressed in most malignant cells. Similar to the HCC dataset, we selected only primary hepatocytes from all other cells found in healthy liver samples. We annotated normal vital organ single cells according to a 3-level cell type classification system (Supplementary Table S5, Fig 4C showing level 1 visualization), similar to the methodology in the human cell atlas [24]. Whenever possible, we used published classification results; in other cases, we used the scPred-based supervised classifier [28].

The co-expression of prominent markers was checked across individual single cells. To do this, we binarized the expression of every marker in each cell. An expression was marked as 1 if the expression surpassed a threshold (indicating presence) else 0 (indicating absence). This led to the creation of a Boolean matrix [m x n], where ‘m’ represents cells and ‘n’ denotes genes. Consequently, for any given gene pair in a specific cell, there are three potential outcomes: 0 (Neither of the genes is expressed in the cell), 1 (One of the genes is expressed in the cell), and 2 (Both genes are expressed in the cell). The significance of co-expression was decided based on p-values derived from the chi-square test with the False Discovery Rate (FDR) correction.

#### Visualization of co-expression pattern using spatial transcriptomic dataset

To further validate the co-expression of top bsAbs pair, we used the filtered ST data from adjacent normal (control), tumor and leading edge (the area between the tumor and the adjacent normal). We used the R package STUtility [29,30] to import all the data followed by integration of the expression data from different sections of each patient using harmony [22]. The Seurat package [31] was used to perform downstream analysis and visualization.

## RESULTS

### Selection of bsAbs pairs based on surface marker expression in bulk RNA-seq

A recent study proposed bi-specific antibodies and tri-specific antibodies (recognizing 2 and 3 antigens) using large bulk RNA-seq databases via Boolean logic [7]. The idea was to identify the markers that separate tumor samples from healthy samples with maximum heterogeneity. However, this study did not account for the angle between markers or capture the varied expression patterns across cell populations. Its selection of the bsAbs was simply determined by the ‘AND’ and ‘NOT’ logic. Specifically, it was based on the localization of the case cluster in the upper-right corner, where every antigen in the pair would have high expression, exemplified by (A AND B), while pairs with other case placements like (A NOT B), (NOT A AND B) or (NOT A NOT B) were disregarded. Nevertheless, it is possible to identify potential bispecific antigens using more robust methods, coupled with a conventional approach for the quantification of sensitivity and specificity.

Our algorithm tends to identify gene pairs that have the maximum difference between case vs. control as well as case vs. healthy vital organs (Fig. 1 A, B). The 7 million gene pairs generated by pairwise combination of 3806 unique genes of the COMPARTMENT database were first screened by the proposed approach (Methods) to assess how well they differentiate between case (HCC samples) and control (adjacent normal liver samples and vital organs). It resulted in over 300 pairs of genes that were significant in terms of both score threshold and Boolean logic: both antigens should have a greater expression in the case cluster (A & B), while the control cluster should display double negative expression patterns (NOT A NOT B) (Fig. 2D, S4). The analysis of the frequency of the genes in bispecific pairs showed a consistent presence of GPC3 across most pairs, followed by PLVAP, CD4, PIGY, and MUC13 (Fig. 2C). Considering the prevalence of GPC3 as a biomarker and therapeutic target in HCC [32,33], its presence in the top most candidate genes was anticipated, confirming the validity of our approach. However, to ensure the specificity of markers for minimizing therapy-related toxicity, the expression of most favorable HCC marker genes was checked in HCC samples as well as vital organs (Fig. 2E). For example, even though PLVAP∼GPC3 showed the best performance in HCC, PLVAP had limited specificity as its comparable levels detected in the lung, kidney, and heart (Fig. 2 A, E). Also, in the CD34 ∼ GPC3 pair, CD34 is specific for hematopoietic stem and progenitor cells [34] suggesting the chance of identifying false positive pairs using only bulk RNA-seq (Supplementary Table S6). Remarkably, among all the genes in these predicted pairs, only MUC13 was specifically expressed in HCC [35].

**Figure 2:**
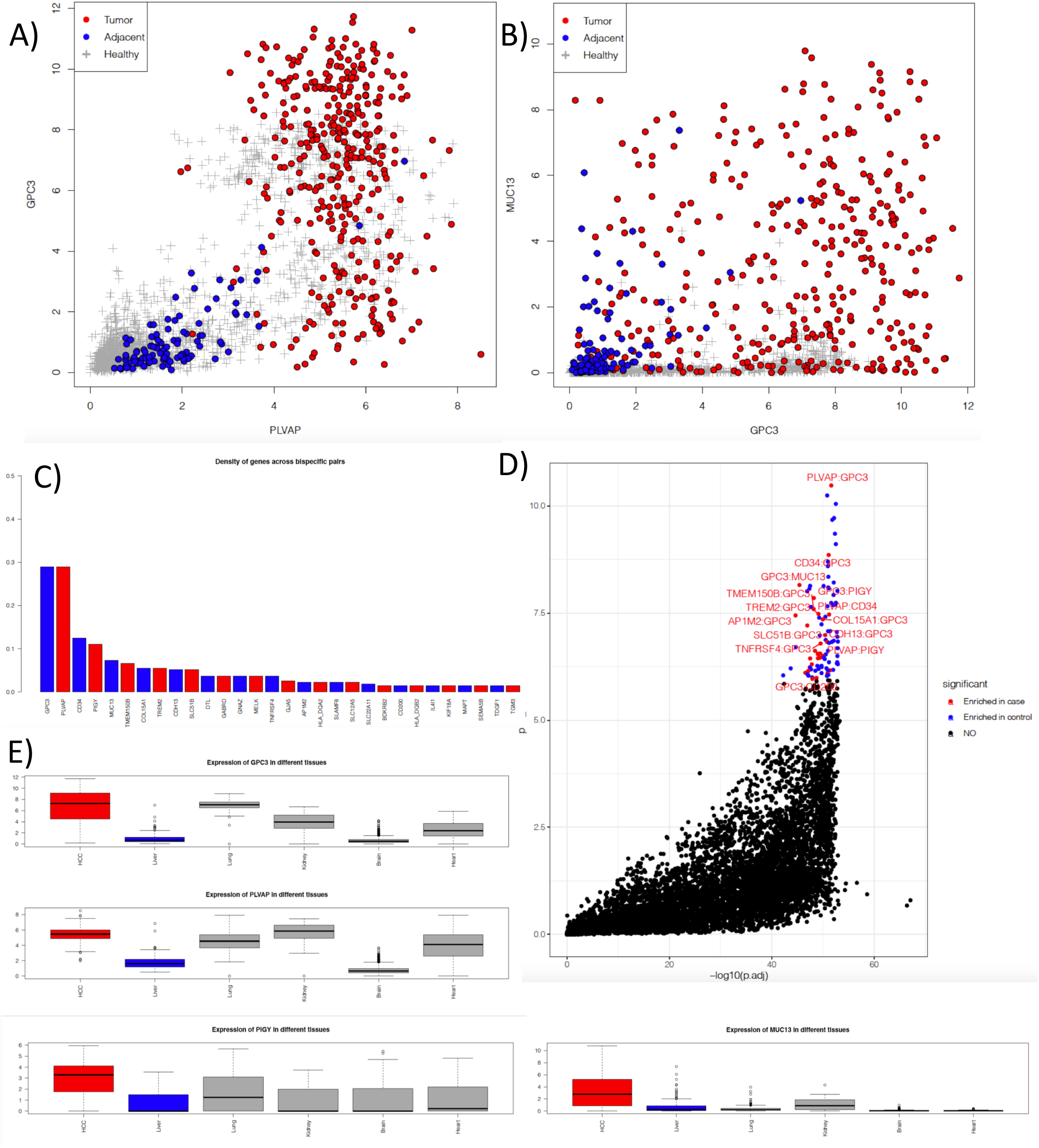
Selection of top bsAbs pairs using OCTAD bulk RNA-seq. **A, B**: Expression distribution of potential marker pairs GPC3∼PLVAP (A) and GPC∼MUC13 (B) in tumors (red), adjacent normal tissues (blue), and normal vital organs (grey). **C**: Frequencies of the most common potential markers for the bsABs candidates. **D**: Score and significance distribution of the proposed bsAbs pairs based on the pair score and permutation-obtained significance scores. **E**: Expression of the four topmost frequent genes of the bsAbs pairs in HCC, normal liver and normal vital organs.

The ability of top candidate marker pairs to separate between tumors vs adjacent normal tissue and vital organs was further assessed by comparing the sensitivity and specificity of SVM models. The mean delta between the sensitivity and specificity for bsAbs pairs and individual markers in the pairs is shown in Fig 3. While we noted a minor increase in sensitivity for the tumor vs. adjacent tissue comparison, a considerable boost in specificity was observed for same comparison (Fig. 3A, Supplementary Table S7). Moreover, both sensitivity and specificity saw marked improvement for comparison between tumors and vital organs (Fig. 3B, Supplementary Table S8). This emphasizes the superiority of dual targeting over single agent targeting.

**Figure 3:**
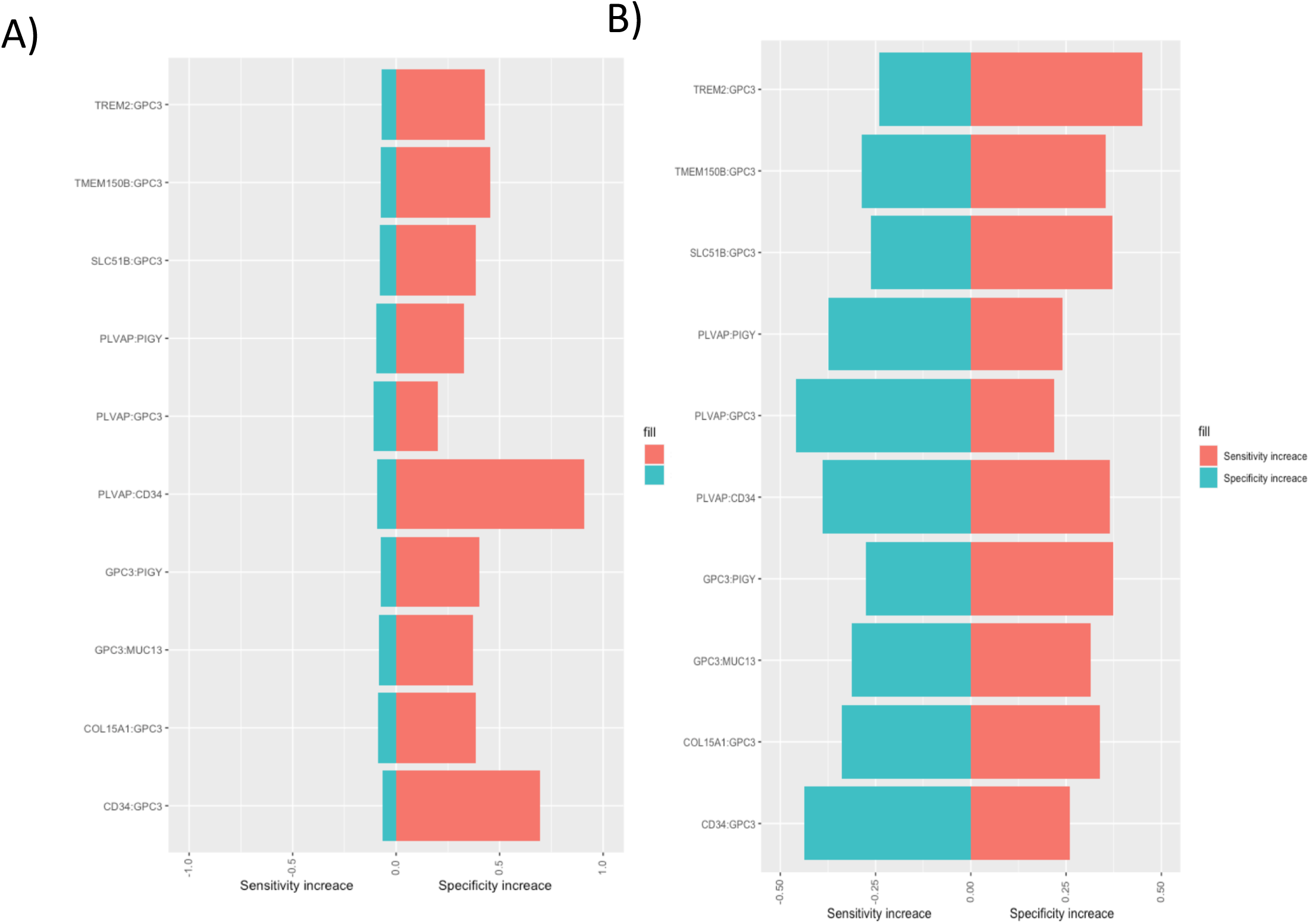
Sensitivity and specificity analysis between dual targeting and single agent targeting from bulk RNA-seq data. Increase of sensitivity and specificity between bsAbs and single agents while using adjacent normal tissues (A) or vital organs (B) as the control.

**Figure 4:**
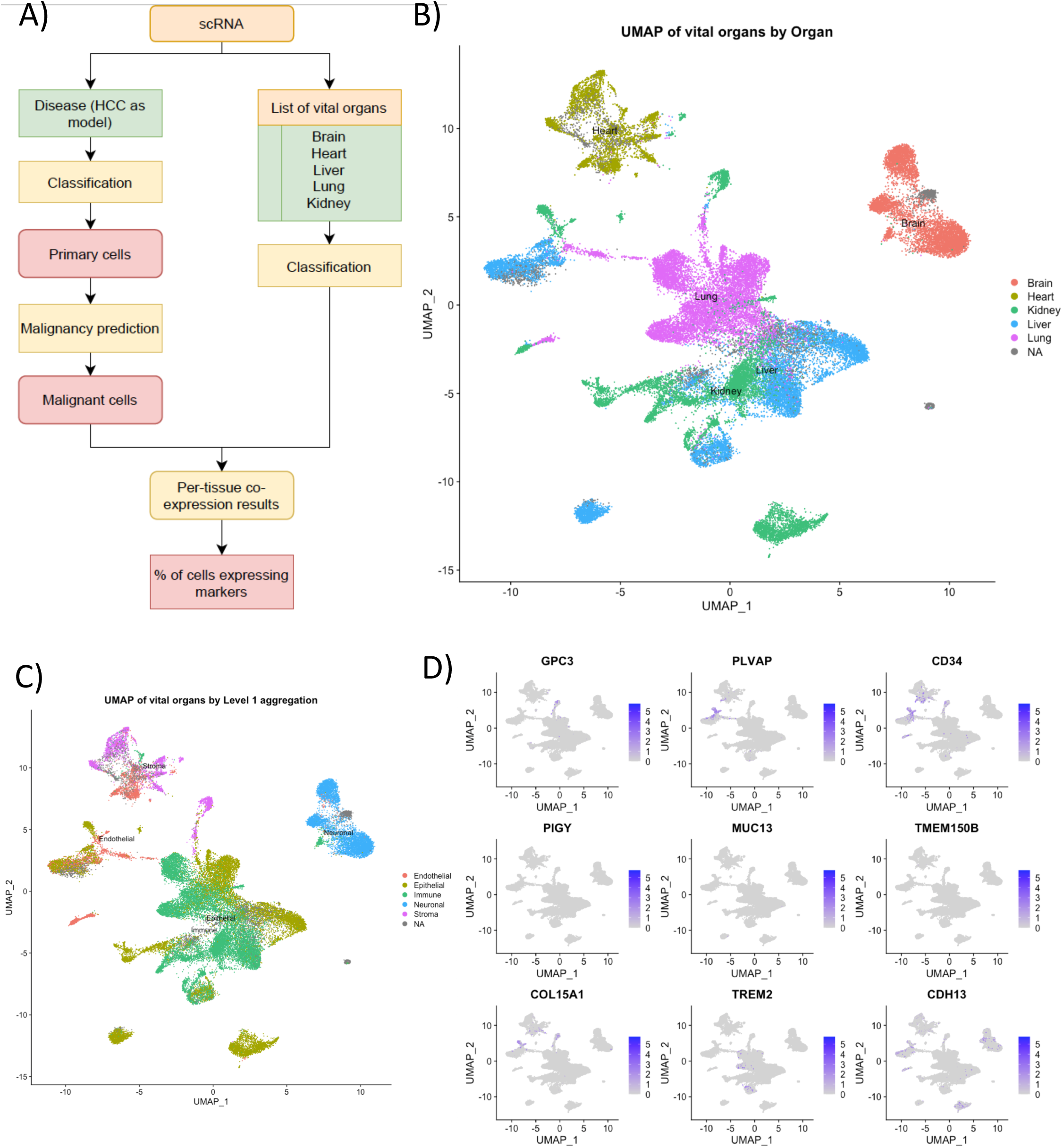
**A**: Principal scheme of the scRNA-seq-based bsAbs step verification. **B**: UMAP of the vital organs scRNA database used for the verification of the cytotoxicity results. **C**: UMAP of the level 1 aggregation to define endothelial, epithelial, immune, neuronal and stromal cells in the vital organs database. **D**: Expression of the top-9 most frequent marker genes across vital organs.

### scRNA-seq quantifies cell-specific co-expression of the bsAbs

Our findings prompted us to leverage the scRNA-seq for the validation of bsAbs derived from bulk RNA-seq, aiming for the discovery of bsAbs with high tissue-specificity and minimal non-specific cytotoxicity. To check the expression of promising marker genes in scRNA dataset, we followed the similar approach as in bulk RNA-seq (Fig. 4A). To evaluate the non-specific cytotoxicity, we quantified the expression level of selected markers on cell surfaces of the vital organs (Fig. 4D). Note that even if the selected pair showed expression in the vital organs, it might remain viable if not expressed in the parenchymal cells of those vital organs.

For the identification of highly sensitive pairs, it is important to concentrate only on the malignant hepatocytes in HCC scRNA data while computing true co-expression percentage. We classified cells, extracted hepatocytes, and identified malignant cells among them. By establishing a threshold derived from the comparison with normal reference hepatocytes (Fig. 5B), out of the 7,140 hepatocytes identified, 5,271 were malignant, and 1,869 were non-malignant (Fig. 5C). Notably, these malignant hepatic cells were classified into five major subtypes, each displaying distinct transcriptional features. This categorization highlighted the inherent heterogeneity within HCC (Fig. 5 C,D).

**Figure 5:**
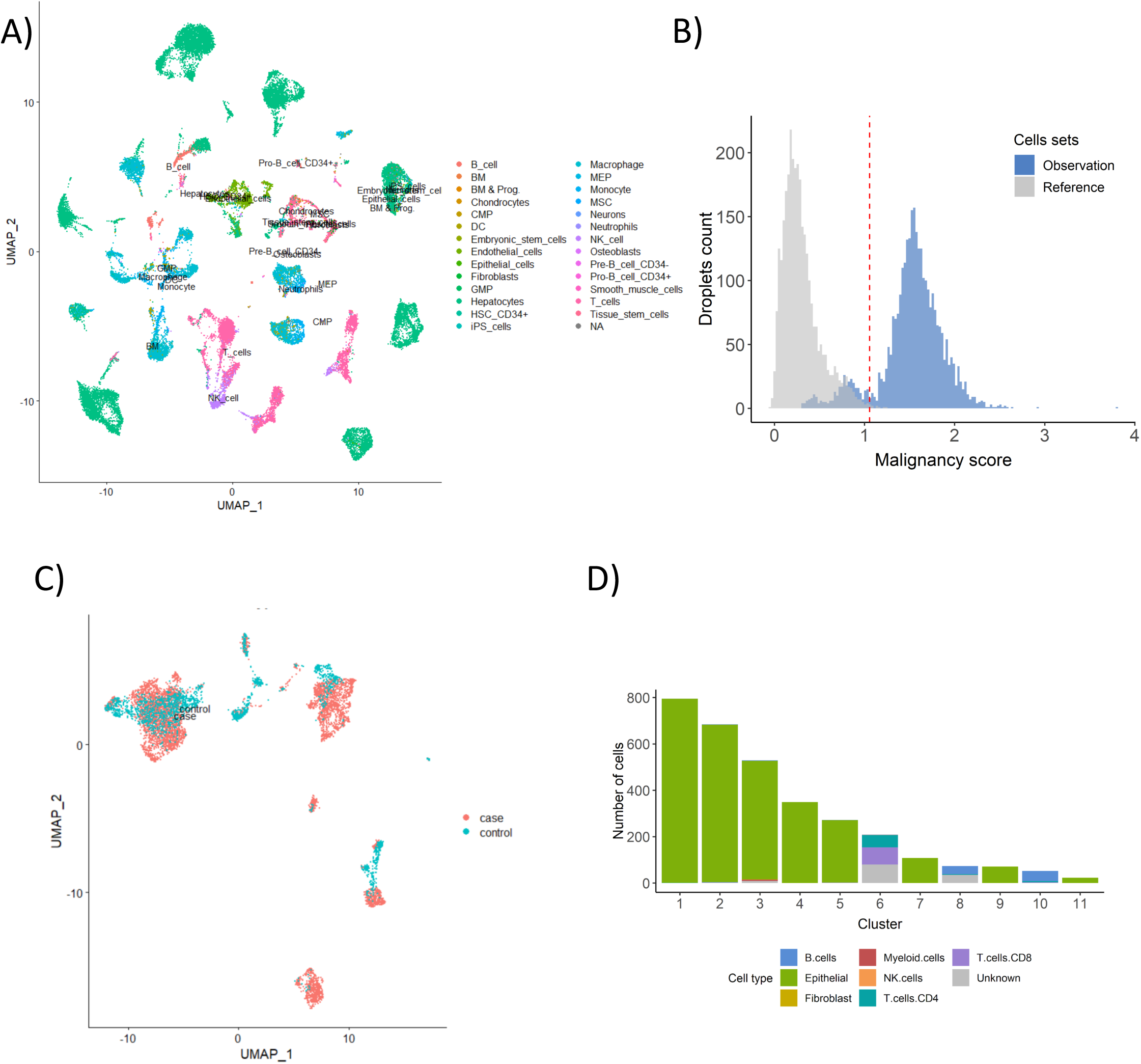
Tumor cell classification and cell malignancy inference from HCC scRNA-seq dataset. **A:** HCC tumor cell composition. **B**: Malignance score distribution of the reference liver dataset vs hepatocytes from the tumor data to leverage only malignant hepatocytes from the tumor data for downstream analysis. **C**: Distribution of the malignant (case) and non-malignant (control) hepatocytes from the HCC scRNA-seq dataset**. D**: The cell count distribution of major cell types found in the HCC scRNA-seq dataset.

The last phase aimed to verify the tissue specificity of the favorable markers at single-cell levels. We first transformed the markers into binary matrices, assigning 1 to a cell if the count surpassed a threshold and 0 otherwise. This binary transformation allowed us to compute the co-expression of the markers across every identified cell type. In HCC, the most promising marker pair turned out to be GPC3∼MUC13. Both markers of this pair are co-expressed in 30% of malignant cells and were barely detected in vital organs. Such a profile strongly suggests the potential of this pair as a promising bsAbs candidate for the HCC therapy (Fig. 6A, B).

**Figure 6:**
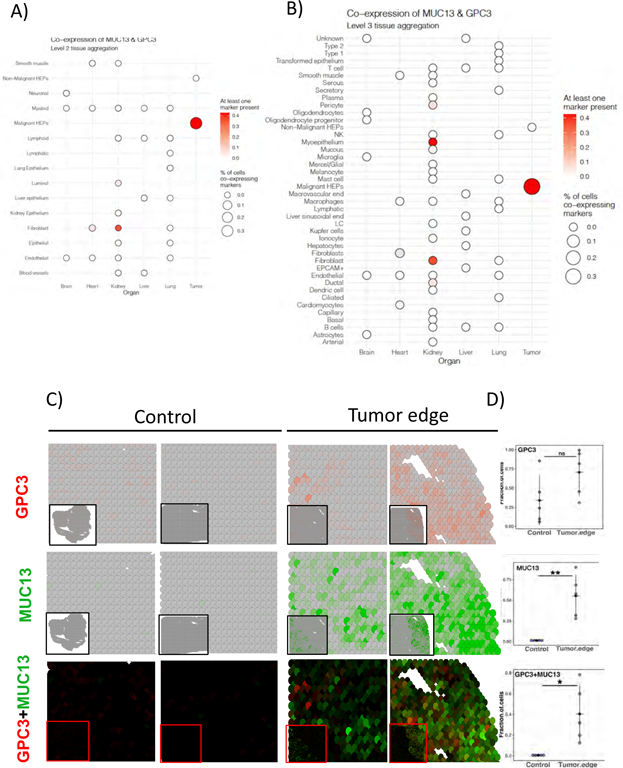
Singe cell and Spatial Transcriptome analysis both confirms the potential of GPC3∼MUC13 marker pair as HCC bsAbs target. **A**: single cell co-expression analysis of the pair on the level II aggregation, **B**: single cell co-expression of the pair in vital organs and malignant hepatic cells on the level III aggregation. C: Expression and co-expression of GPC3 and MUC13 in control and tumor spatial transcriptome D: Percentage of cells expressing and co-expressing GPC3 and MUC13 in spatial datasets

### Validation of specificity and sensitivity of topmost bsAbs pair using spatial transcriptomic data

The spatial transcriptomic analysis unveiled a distinctive pattern in the tumor microenvironment, where both GPC3 and MUC13 were highly expressed in tumor samples contrasting sharply with their complete absence in the control sample (Fig. 6C and supplementary figure 1). GPC3 and MUC13 were co-expressed in approximately 40% of tumor cells (Fig. 6D and supplementary figure 1).

## DISCUSSION AND CONCLUSION

Our newly developed computational pipeline underscores the strength of an integrated approach in the unbiased survey of bsAbs candidates for a cancer. By fusing the complementary advantages of both bulk and single-cell RNA sequencing, we can accentuate their positive attributes while simultaneously offsetting the limitations inherent when each is employed in isolation. Examining the spatial distribution of these markers within tumors proved valuable while assessing their potential in bsAbs development. Even though our current study is anchored around HCC, the flexibility and adaptability of our pipeline mean that it can be swiftly recalibrated to identify bsAbs for a diverse range of cancers. One salient insight from our HCC case study was the discernible value added by scRNA-seq in curtailing the chances of discovering false positive pairs derived from bulk RNA-seq. The visualization of the top pair GPC3∼MUC13 *in situ* further provides evidence for their co-expression in tumor cells, suggesting their potential as a therapeutic target. However, given the innate data dropout challenge with scRNA-seq and the potential discordance between RNA and actual protein expression, it is imperative that the promising marker pairs identified through our pipeline should undergo subsequent validation. Techniques such as immunostaining or other protein quantification methods would be instrumental in this confirmatory phase.

## Code and data availability

All bulk data used in the paper is available online: OCTAD database (https://github.com/Bin-Chen-Lab/octad). Also, standalone R package for bsAbs testing in bulk data are available via GitHub (https://github.com/Lionir/bsAbsFinder).

## Acknowledgement

This work is partially supported by Hengenix Biotech, Inc. We thank the members from the Hengenix Biotech and Chen Lab for the fruitful discussion.

## Author Contributions

E.C. and B.C. conceived the study. E.C performed all the computational analysis with the support of R.S. S.P., W.X, and J.X.. S.P. led the submission and revision. E.C., S.P. and B.C. wrote the manuscript. B.C. supervised the study.

---

**Supplementary Fig 1.**
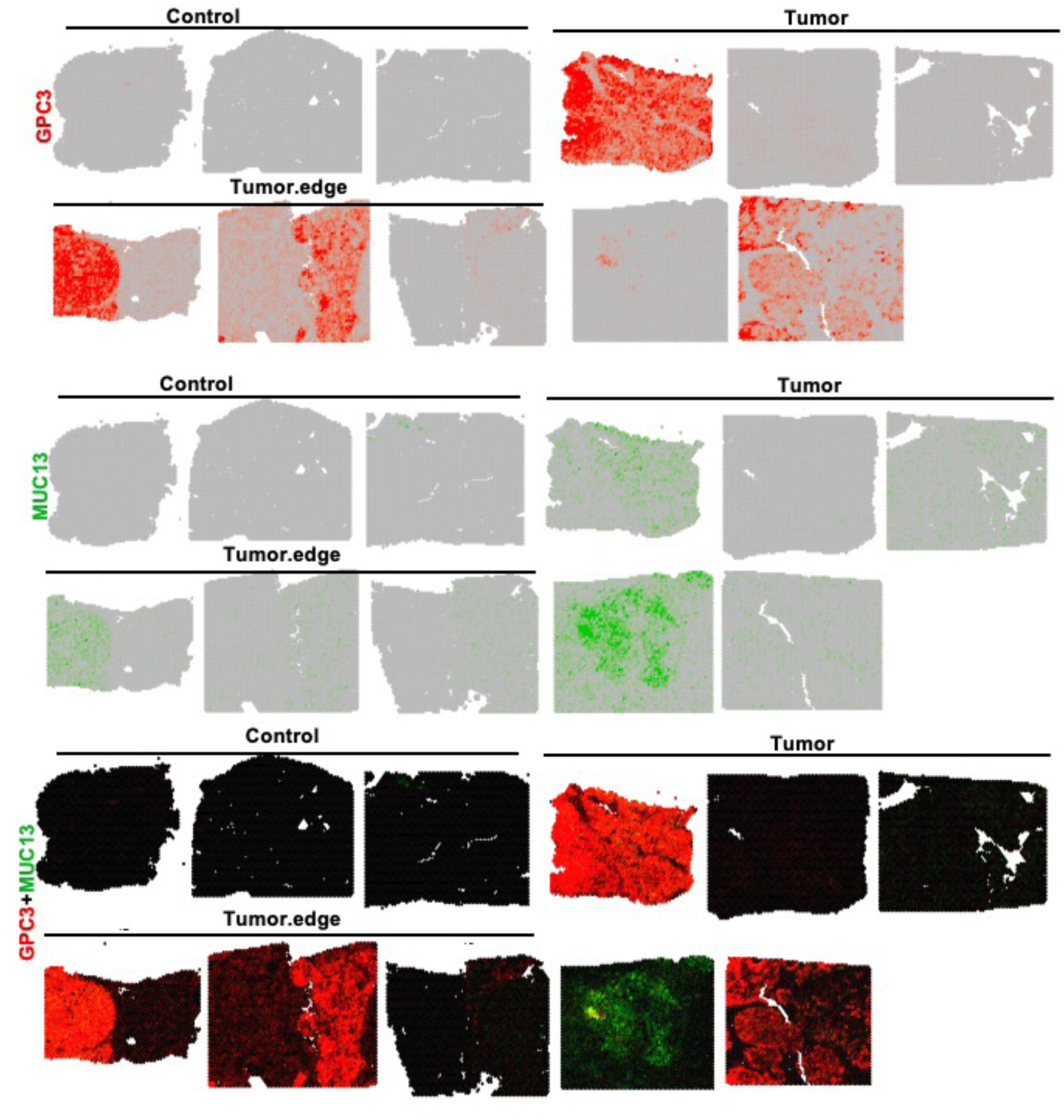
Additional Spatial Transcriptome Slides

**Supplementary table 1:** RNA-seq samples of hepatocellular carcinoma from the OCTAD were included in the study.

**Supplementary table 2:** List of RNA-seq samples from vital organs (Liver, Lung, Kidney, Heart, Brain) obtained from the OCTAD database to validate the expression of potential pairs in vital organs

**Supplementary table 3:** List of surface-expressed proteins derived from the COMPARTMENT database used in the study.

**Supplementary table 4:** List of single-cell RNA-seq experiments included in the study.

**Supplementary table 5:** List of potential genes significantly co-expressing on the surface of tumor cells in comparison with adjacent tissue and healthy vital organs in bulk RNA-seq.

**Supplementary table 6:** 3 levels of cell populations in scRNA-seq used for the verification of candidates for bsAbs targeting.

**Supplementary table 7:** Specificity and sensitivity comparison between single markers and each antigen with the relative power to distinct tumor samples from a healthy liver. The specificity and sensitivity column indicates corresponding values for marker pairs, while specificity and sensitivity for marker 1 and marker 2 indicate corresponding values for individual genes to distinct tumor samples from healthy liver samples.

**Supplementary table 8:** Specificity and sensitivity comparison between single markers and each antigen with the relative power to distinct tumor samples from healthy vital organs. The sensitivity and specificity columns indicate corresponding values for maker pairs, while specificity and sensitivity for marker 1 and marker 2 indicate corresponding values for individual genes to distinct tumor samples from healthy vital organ samples.

## References

1. Huang S, Van Duijnhoven SMJ, Sijts AJAM et al. Bispecific antibodies targeting dual tumor-associated antigens in cancer therapy. J Cancer Res Clin Oncol 2020;146:3111–22.

2. Sellmann C, Doerner A, Knuehl C et al. Balancing Selectivity and Efficacy of Bispecific Epidermal Growth Factor Receptor (EGFR) × c-MET Antibodies and Antibody-Drug Conjugates*. Journal of Biological Chemistry 2016;291:25106–19.

3. Spiegel JY, Patel S, Muffly L et al. CAR T cells with dual targeting of CD19 and CD22 in adult patients with recurrent or refractory B cell malignancies: a phase 1 trial. Nat Med 2021;27:1419–31.

4. Hirabayashi K, Du H, Xu Y et al. Dual-targeting CAR-T cells with optimal co-stimulation and metabolic fitness enhance antitumor activity and prevent escape in solid tumors. Nat Cancer 2021;2:904–18.

5. Lajoie MJ, Boyken SE, Salter AI et al. Designed protein logic to target cells with precise combinations of surface antigens. Science 2020;369:1637–43.

6. Oostindie SC, Rinaldi DA, Zom GG et al. Logic-gated antibody pairs that selectively act on cells co-expressing two antigens. Nat Biotechnol 2022;40:1509–19.

7. Dannenfelser R, Allen GM, VanderSluis B et al. Discriminatory Power of Combinatorial Antigen Recognition in Cancer T Cell Therapies. Cell Systems 2020;11:215–228.e5.

8. Robinson DR, Wu Y-M, Lonigro RJ et al. Integrative clinical genomics of metastatic cancer. Nature 2017;548:297–303.

9. Zeng WZD, Glicksberg BS, Li Y et al. Selecting precise reference normal tissue samples for cancer research using a deep learning approach. BMC Medical Genomics 2019;12:21.

10. Zeng B, Glicksberg BS, Newbury P et al. OCTAD: an open workspace for virtually screening therapeutics targeting precise cancer patient groups using gene expression features. Nat Protoc 2021;16:728–53.

11. Misek SA, Newbury PA, Chekalin E et al. Ibrutinib Blocks YAP1 Activation and Reverses BRAF Inhibitor Resistance in Melanoma Cells. Mol Pharmacol 2022;101:1–12.

12. Zhao G, Newbury P, Ishi Y et al. Reversal of cancer gene expression identifies repurposed drugs for diffuse intrinsic pontine glioma. Acta Neuropathologica Communications 2022;10:150.

13. Binder JX, Pletscher-Frankild S, Tsafou K et al. COMPARTMENTS: unification and visualization of protein subcellular localization evidence. Database 2014;2014:bau012.

14. Lu Y, Yang A, Quan C et al. A single-cell atlas of the multicellular ecosystem of primary and metastatic hepatocellular carcinoma. Nat Commun 2022;13:4594.

15. Aizarani N, Saviano A, Sagar et al. A human liver cell atlas reveals heterogeneity and epithelial progenitors. Nature 2019;572:199–204.

16. Vieira Braga FA, Kar G, Berg M et al. A cellular census of human lungs identifies novel cell states in health and in asthma. Nat Med 2019;25:1153–63.

17. Ferreira RM, Sabo AR, Winfree S et al. Integration of spatial and single-cell transcriptomics localizes epithelial cell–immune cross-talk in kidney injury. JCI Insight 2021;6, DOI: 10.1172/jci.insight.147703.

18. Habib N, Avraham-Davidi I, Basu A et al. Massively parallel single-nucleus RNA-seq with DroNc-seq. Nat Methods 2017;14:955–8.

19. Wang L, Yu P, Zhou B et al. Single-cell reconstruction of the adult human heart during heart failure and recovery reveals the cellular landscape underlying cardiac function. Nat Cell Biol 2020;22:108–19.

20. Satija R, Farrell JA, Gennert D et al. Spatial reconstruction of single-cell gene expression data. Nat Biotechnol 2015;33:495–502.

21. Trapnell C, Cacchiarelli D, Grimsby J et al. The dynamics and regulators of cell fate decisions are revealed by pseudotemporal ordering of single cells. Nat Biotechnol 2014;32:381– 6.

22. Korsunsky I, Millard N, Fan J et al. Fast, sensitive and accurate integration of single-cell data with Harmony. Nat Methods 2019;16:1289–96.

23. Aran D, Looney AP, Liu L et al. Reference-based analysis of lung single-cell sequencing reveals a transitional profibrotic macrophage. Nat Immunol 2019;20:163–72.

24. Regev A, Teichmann SA, Lander ES et al. The Human Cell Atlas. Gingeras TR (ed.). eLife 2017;6:e27041.

25. Wu R, Guo W, Qiu X et al. Comprehensive analysis of spatial architecture in primary liver cancer. Science Advances 2021;7:eabg3750.

26. broadinstitute/infercnv. 2023.

27. Visualizing Large-scale Copy Number Variation in Single-Cell RNA-Seq Expression Data.

28. Alquicira-Hernandez J, Sathe A, Ji HP et al. scPred: accurate supervised method for cell-type classification from single-cell RNA-seq data. Genome Biology 2019;20:264.

29. ludvigla. ludvigla/STUtility. 2019.

30. Bergenstråhle J, Larsson L, Lundeberg J. Seamless integration of image and molecular analysis for spatial transcriptomics workflows. BMC Genomics 2020;21:482.

31. Stuart T, Butler A, Hoffman P et al. Comprehensive Integration of Single-Cell Data. Cell 2019;177:1888–1902.e21.

32. Koksal AR, Thevenot P, Aydin Y et al. Impaired Autophagy Response in Hepatocellular Carcinomas Enriches Glypican-3 in Exosomes, Not in the Microvesicles. JHC 2022;9:959–72.

33. Wu T, Song Z, Huang H et al. Construction and evaluation of GPC3-targeted immunotoxins as a novel therapeutic modality for hepatocellular carcinoma. International Immunopharmacology 2022;113:109393.

34. Sidney LE, Branch MJ, Dunphy SE et al. Concise Review: Evidence for CD34 as a Common Marker for Diverse Progenitors. Stem Cells 2014;32:1380–9.

35. Dai Y, Liu L, Zeng T et al. Overexpression of MUC13, a Poor Prognostic Predictor, Promotes Cell Growth by Activating Wnt Signaling in Hepatocellular Carcinoma. Am J Pathol 2018;188:378–91.

